# Seasonal frost improves probiotic and nutrient availability in fermented vegetables

**DOI:** 10.1101/2025.06.27.661941

**Authors:** Andrew Luzmore, Jason Grauer, Dan Barber, Grace Jorgensen, Pearson Lau, Swapan Jain, Gabriel G. Perron

**Author notes:** These authors contributed equally to this work (Andrew Luzmore, Jason Grauer). Corresponding authors: Department of Biology Reem-Kayden Center for Science and Computation Bard College 30 Campus Road, Annandale-on-Hudson, NY, 12571.

## Abstract

Climate change is shifting seasonal patterns in temperate regions, increasing the likelihood of early frost and raising questions about its impact on food production and quality. In this study, we tested how a single seasonal frost event influences microbial communities, fermentation dynamics, and the nutritional profile of naturally fermented cabbage and carrots, two cold-tolerant crops commonly grown in the Northeast US Using microbial sequencing, metagenomic analysis, and targeted vitamin assays, we found that frost exposure had no negative impact on microbial diversity or fermentation outcomes. Instead, it was associated with subtle shifts in microbial composition and increased abundance of genes involved in vitamin biosynthesis, including those linked to vitamin K□, B□□, and threonine. These genetic changes corresponded to higher concentrations of vitamins A and E in fermented carrots and vitamin K□ in fermented cabbage. Our findings suggest that frost can enhance the nutritional and sensory qualities of fermented vegetables, offering a model for a new climate-resilient strategy for producing value-added, regionally distinctive food products.

## 1. Introduction

Global food systems focused on production exert considerable financial, human, and environmental pressures on producers and consumers worldwide ^1^. The drive to scale up production for competitive advantage ^2^ has led to geopolitical constraints shaping hubs of food production ^3,4^, often resulting in non-native crops dominating local and global economies ^5^. Concentrated agriculture in these hubs requires extensive inputs and modifications, which, while improving production costs in specific areas, have significant impacts on environmental health, biodiversity loss ^6,7^, and climate change ^8^.

Large-scale agricultural practices have a significant impact on global freshwater resources. For instance, agriculture uses approximately 80% of California’s developed water resources ^9^, only exacerbating water scarcity during droughts, impacting drinking water availability ^10^, and leading to conflicts with other sectors ^11^. Interestingly, the state leads the US in cabbage (22.9 million cwt) and carrot (6.7 million cwt) production ^12^, even though the two crops were historically bred in the cool, humid climates of coastal and northern Europe ^13,14^. Producing cabbage in California’s arid climate requires between 326,000 and 489,000 gallons of water per acre ^15^, amounting to 2-7 times greater than in regions with cooler temperatures and higher precipitation, such as New York State, where rainfall supplies the majority of water ^16^. Beyond water use, intensified agricultural practices in centralized production hubs also contribute to land degradation, particularly through monocropping, which depletes soil nutrients, reduces organic matter, and exacerbates soil erosion ^17,18^. Such degradation diminishes soil fertility, necessitating greater application of synthetic nitrogen fertilizers to sustain yields, which in turn amplifies greenhouse gas emissions, notably nitrous oxide as well as CO□ and CH□, thereby contributing to global climate change ^19,20^.

Over the past few decades, the American Northeast, characterized by lower temperatures, shorter day lengths, and distinct seasonal changes in the fall, has observed changes in seasonal patterns that are mainly explained by a delay in the first autumn freeze ^21,22^. Such changes also increased the unpredictability of rapid temperature fluctuations ^23^, increasing the risk of exposure to cold temperatures and frost, which can affect the biochemical and physiological properties of fruits and vegetables ^24^. Frost, for example, causes physical and biochemical changes in vegetables as freezing water bursts the cell walls, altering the cell content and enzymatic activities while increasing oxidative stress and increasing frost injuries due to pathogen infection ^25^. The latter can thus have important implications for plant growth, food safety, downstream application in cooking, and ultimately, market value.

In response to these environmental changes, producers are employing various strategies to enhance resilience in the face of climate change and to extend their farming seasons both on the farm and at the market ^26^. For example, heated and semi-heated greenhouses can be used to artificially create ideal atmospheric conditions for seed starting or year-round production ^27^. Similarly, low- and high-tunnel structures can be used to protect crops from frost and extend the growing season by several months ^28,29^. Another approach is to use cold-tolerant crops that can be harvested later in the fall or that overwinter in the soil and are harvested in the spring when growth resumes ^30,31^. Allowing cold-tolerant crops such as carrots and cabbage to remain in the soil is known to increase the plant’s sweetness ^32^ and subsequent storability in cold rooms ^33^. Cabbages produce phytochemicals such as glucosinolates in response to cold, which is likely responsible for the plant’s increased storability and possibly nutritional quality ^34^.

Considering the advantages of using cold-tolerant crops, we investigated whether frost exposure at harvest alters microbial succession and nutrient profiles in fermented cabbage and carrots, potentially enhancing sensory and nutritional qualities. Fermentation, a traditional preservation technique that has regained popularity ^35^, is another strategy used by farmers to extend their crop’s presence at the market while retaining a sense of freshness beyond the growing season ^36,37^. To do so, we tested the effect of a single 12-hour frost event at harvest on the probiotic diversity and nutritional quality of fermented cabbages, specifically sauerkraut, and carrots by comparing individual plants exposed to frost with those protected from frost using agricultural tunnels. The study was conducted in collaboration with the farming staff at Stone Barns Center for Agriculture and Food (Pocantico Hills, NY), a leading research center for regenerative agriculture, and the chefs at Blue Hill at Stone Barns (Pocantico Hills, NY), a leading restaurant recognized worldwide for its innovative approach to local food systems and sustainability.

In brief, our study demonstrates that the use of cold-tolerant crops as a season extension strategy produces ferments with higher probiotic and nutritional properties, and enhances flavor complexity in two commercially important vegetables, both regionally and globally. In addition, our study contributes to the development of a more sustainable and local economy in the Northeast by establishing a greater potential for premium market positioning of products unique to the region’s climate.

## 2. Materials and methods

### 2.1 Study Sites & Growing Conditions

The study was conducted at the Stone Barns Center for Food & Agriculture in Pocantico Hills, NY, situated on 500 acres of both private and public land (**Figure 1**). As part of a functioning farming operation, all crops used in this experiment were part of the regular farming crop schedule from seeding to harvest. To this end, ‘Murdoc’ cabbage (*Brassica oleracea* var. *capitata*), a hybrid variety known for its large, pointed heads and a typical weight of 3 to 4 Kg at maturity and often used in sauerkraut making, were seeded in our greenhouse on June 30, 2023. We then transplanted the starter cabbages into the main vegetable field on July 24, 2023, where they remained until harvest. We also direct-seeded ‘Bolero’ carrot variety (*Daucus carota* subsp. *sativus*), characterized by a traditional blunt end that reaches about 20 cm at maturity, in the main vegetable field on July 28, 2023, where they remained until harvest. Weather conditions for the growing season (i.e., July-October) were monitored daily using the closest weather station, Harrison, NY, available via wundergrounds.com. While July 2023 was marginally cooler and drier when compared to July’s ten-year average (i.e., 2015-2024), weather conditions over the growing season did not differ markedly from the annual average for the same growing season (Table S1). To this end, organic lime, potash, and pulverized kelp were added at the beginning of the season. Compost and crab meal were added before transplanting the cabbages as well. Finally, a drip irrigation system provided 1 inch of water per week.

**Figure 1.**
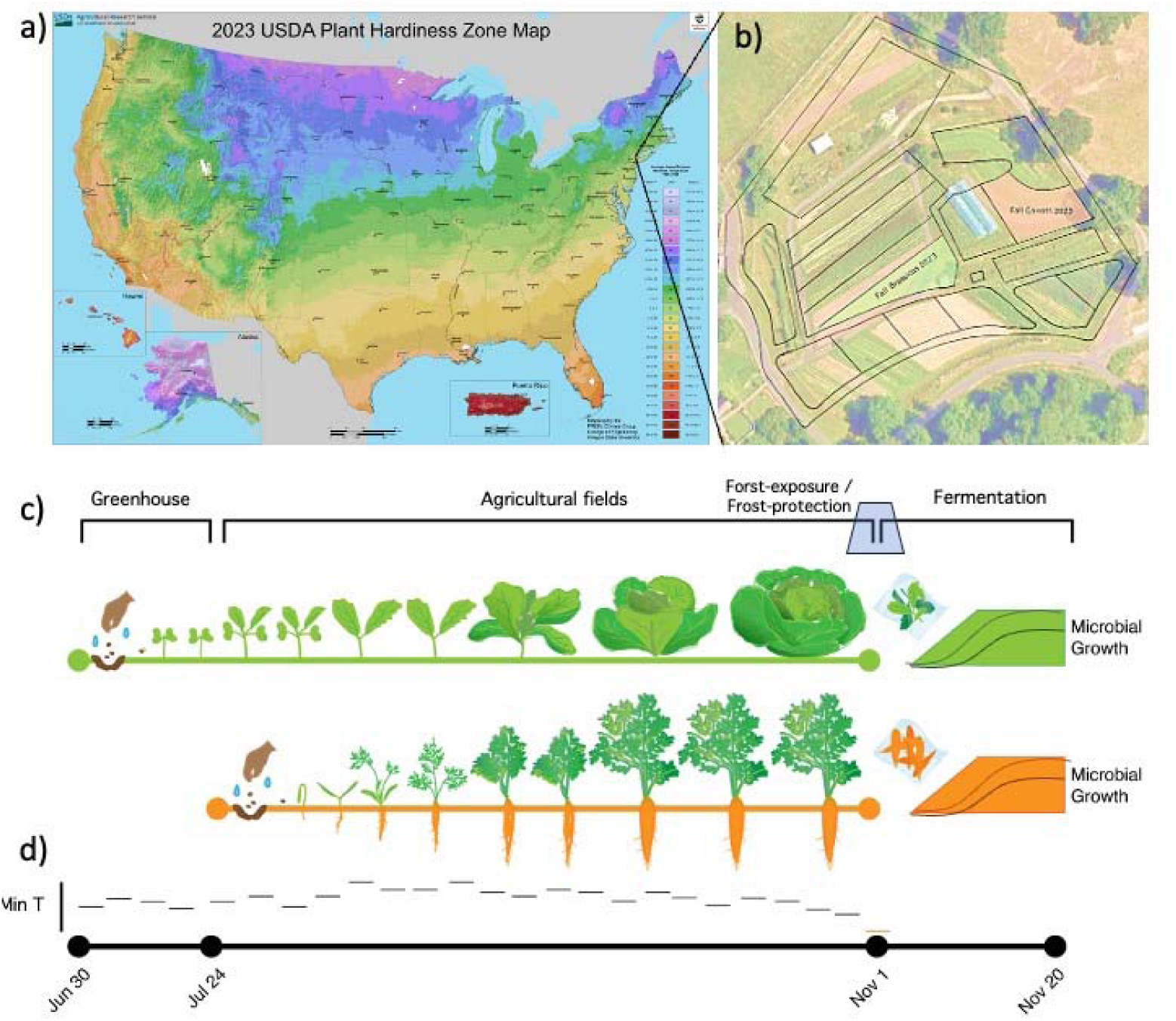
Experimental Design Investigating the Impact of Frost on Fermented Vgetables. **a**) The study was conducted in Pocantico Hills (NY), which is characterized by cold climates in the fall compared to other climate zones in the USA. The map shows the USDA Plant Hardiness Zone (https://planthardiness.ars.usda.gov/pages/map-downloads). **b**) Study compared three individual carrots and cabbages grown in separate fields at the Stone Barns Center for Food and Agriculture, Pocantico Hill (NY, and **c**) compared microbial probiotic diversity and nutritional quality between ferments made from frost-conditioned carrots and cabbages to ferments made from vegetables protected from frost exposure. **D**) The first seasonal frost happened in the early morning of November 1, 2023, following an average growing season that started June 30, 2023, for cabbages and July 24, 2023. Samples from the ferment were taken after the fermentation was completed on November 20, 2023.

### 2.2 Season frost exposure treatments

All cabbages and carrots used in this study were grown with minimal disturbance until the first seasonal frost, defined as air temperature remaining below < 0□ for at least one hour. Crucially, to test the effect of the first seasonal frost, we protected half of the crops with season extension devices and left the other half exposed to ambient temperatures. For cabbages, we set up a ‘low tunnel’ made of a combination of bent PVC pipe hoops with polyethylene greenhouse plastic covers (Tufflite IV™, Berry Global, Evansville, IN). For carrots, we used a double layer of ‘reemay’ row cover fabric (Abrigon™, JMI Legacy Manufacturing, Ponchatoula, LA) to protect half of the carrots, leaving the other half exposed at all times. To minimize their potential impact on the crop, these covers were installed on half of each treatment area once temperatures were projected to drop to 4.5□ or below at night. Given the more breathable nature of the row cover fabric placed over the carrots, once covered, the carrot row cover was left on throughout the day and night until we encountered frost conditions. The cabbage cover was temporarily removed on days when the sun was brightest and temperatures were predicted to reach above 10□. However, it was always put back on by 4 pm each day and remained in place throughout the evening until the sun set and temperatures dropped. Using four temperature sensors (SensorPush, Brooklyn, NY) located in the middle of each crop, we observed the first seasonal frost during the night of November 1, 2023, when air temperature plummeted below 0□ for 12 hours (Figure S1). The same probes also confirm that the wind tunnel protected the crop with air temperature reaching a minimum of 0.1□ for cabbages and 0.05□ for carrots.

### 2.3 Harvest and Fermentation Conditions

Following the seasonal frost event, we harvested four cabbages from each treatment and four carrots from each treatment at 9 am the following morning. For each cabbage collected, we removed and finely shredded the leaves (approximately 0.1 cm thick) to yield 200 g and added 2% by weight (4 g) of fine sea salt (La Baleine, Provence, France). The cabbage and salt mixture was mashed under sterile conditions for 30 seconds, from which 50 g was transferred to a vacuum bag and sealed at 100% pressure in a cryo-vac machine (Sammic, Ivantson, IL). This procedure was repeated for each cabbage for four independent replicates for each treatment. The samples were incubated in a temperature-controlled room (12.8□) to allow for fermentation until the ferments developed the requisite amount of perceived acidity for use in the restaurant. For the frost-protected cabbage, fermentation visibly slowed down after 23 days, with the absence of carbon dioxide production after touch. Using pH strips (Precision Laboratories, Cottonwood, AZ), we observed that the pH level no longer decreased and reached a final pH of 4.0. For frost-exposed cabbages, fermentation visibly slowed down, and the perceived acidity reached the required level after 35 days. Using pH strips, we observed that the pH level no longer decreased and reached a final pH of 4.25.

For each carrot we picked, the vegetable was washed under running tap water and sliced to a consistent thickness of 0.2 cm using a Japanese mandolin, yielding 200 g. We added 2% by weight (4 g) of fine sea salt (LaBaleine, Provence, France) and let the sliced carrots sit with the salt for five minutes, ensuring the salt was evenly distributed. For each sliced carrot and salt mixture, we then transferred 50 g of it to a sterile vacuum bag and sealed at 100% pressure for a total of four independent replicates. The samples were then transferred to a temperature-controlled room (12.8□) to allow for fermentation. For the frost-protected carrots, fermentation visibly slowed down after 23 days, with the absence of carbon dioxide production after touch. Using pH strips (Precision Laboratories, Cottonwood, AZ), we observed that the pH level no longer decreased and reached a final pH of 4.5. For frost-exposed carrots, fermentation visibly slowed down, and the perceived acidity reached the required level after 35 days. Using pH strips, we observed that the pH level no longer decreased and reached a final pH of 4.75.

### 2.4 Sample collection and processing

For each sample, we collected 1 mL of brine after carefully mixing the ferments in the vacuum bag for 30 seconds, thereby including microbial communities on the leaf surfaces. The brines and the ferments were stored at -80□ until further analysis. We extracted microbial DNA from 150 μL of brine using the procedure described in the MoBio Power Food DNA Extraction kit (MoBio, Carlsbad, CA). To investigate microbial diversity, we first used a *16S rRNA* amplicon sequencing approach to characterize the bacterial communities and the *ITS2* amplicon sequencing approach to characterize the fungal communities found in each ferment. More specifically, samples were prepared for *16S* amplicon sequencing and *ITS2* using a modified version of the Illumina 16S Metagenomics Sequencing Protocol. The V4 region of the *16S rRNA* gene using the Golay-barcoded primers 515F and 806R ^38^. while primers ITS3F and ITS4R were used to amplify the *ITS2* region ^39^. Following SPRI bead cleanup (Beckman Coulter, Brea, CA), libraries were pooled at equimolar ratios and sequenced on the Illumina NextSeq 2000 platform using a 600-cycle flow cell kit to produce 2 × 300 bp paired-end reads (Illumina, San Diego, CA). Finally, we used whole-genome sequencing to describe the total gene content of each ferment. Samples were prepared for whole-genome sequencing using the Illumina DNA Prep tagmentation kit and IDT For Illumina Unique Dual Indexes. Sequencing was performed on the Illumina NextSeq 2000 platform using a 300-cycle flow cell kit to produce 2 × 150 bp paired-end reads. All sample processing and sequencing were conducted by SeqCoast Genomics (Portsmouth, NH). All unprocessed sequence reads are available at the Sequence Read Archive of the National Center for Biotechnology Information (NCBI accession number: PRJNA1281670).

### 2.5 Quantifying microbial diversity

We characterized bacterial diversity from *16S* amplicons and fungal diversity from *ITS2* amplicons for each ferment by identifying and tabulating the number of different amplicon sequence variants (i.e., ASVs). Sequence variants can then be assigned to a taxonomic rank, usually at the genus level, providing additional information about the biology of each microbiome community. More specifically, we processed the *16S rRNA* and *ITS2* reads using the *DADA2* pipeline version 1.26 ^40^ using standard parameters unless specified and implemented in *R* version 4.4.2. In total, we obtained 1,160,000 pairs of forward and reverse reads for *16S rRNA* and 10,863,618 pairs of reads for *ITS2*, with an average read length of 250 base pairs, totaling approximately 583 G bases and an average sequencing depth per sample of 41,642.9 paired reads. Each sequence read was then quality-checked, trimmed (i.e., forward reads to 240 bp and reverse reads to 225 bp), assessed for chimeric contaminants, and de-noised for possible sequencing errors. After processing the reads for quality control, we were able to conserve 980,261 (84.1% of the initial) paired reads.

Taxonomy was assigned using both the DADA2 native taxa identifier function trained on the SILVA ribosomal RNA gene database version 138.1 ^41,42^ for 16S rRNA and the UNITE version 10.0 for ITS2 identification ^43^. A complete list of all ASVs and their abundance in each sample is provided in **Table S2** for *16S rRNA* and in **Table S3** for *ITS2*. Additionally, complete taxonomic assignments are provided in **Tables S4** and **S5** for *16S* and *ITS2,* respectively. Additionally, a mapping file linking sample names to the various treatments is provided in **Table S6**.

### 2.6 Microbial Community Analysis

The diversity of microbiomes was analyzed using *phyloseq* version 1.30.0 ^44^. For both bacterial and fungal datasets, we estimated the number of Observed ASVs and Pielou’s Evenness, on samples rarefied to the lowest sampling depth. The latter includes evenness measures among the different ASVs present in a sample, providing an overall measure of diversity based on relative abundance ^45^. We tested for differences between treatments using linear modeling and comparing the different statistical models with Akaike’s Information Criterion (AIC).

To test whether there were statistical differences in population structure between treatments (e.g., frost-exposed cabbage vs. frost-protected cabbage), we performed Principal Coordinate Analysis (PCoA) on distance matrices estimated from the Bray-Curtis dissimilarity index and phylogenetic signal, calculated as weighted UniFrac distance scores ^46^. We used a permutational multivariate analysis of variance (PERMANOVA), implemented via the *’adonis’* function of the *vegan* package version 2.5.6, to test for significance ^47,47^. The latter is a non-parametric method that estimates *F*-values from distance matrices and relies on permutations to determine the statistical significance of the difference between group means. All computations were performed in *R* version 4.4.2, and data visualization was done using *ggplot2* ^48^.

### 2.7 Metagenomic Analysis

To perform bioinformatic and subsequent whole-genome sequencing, we uploaded our demultiplexed reads onto the KBase platform. Reads were quality-checked with *FastQC* version 0.12.1, filtered with *Trimmomatic* version 0.36, and assembled into contigs using *metaSPAdes* version 3.15.3. Contigs were annotated using the KBase annotation module, which contains components from the Rapid Annotation using Subsystems Technology toolkit (*RASTtk*, version 1.073) ^49^. Outputs from the annotation include not only predictive taxonomy but also functional pathways from the PUBSEED database (version 2.0) ^50^. Gene counts associated with predictive pathways were quantified from the output results of the View Function Profile for Genomes (version 1.4.0) in *R* version 4.4.2.

We then constructed the *mem* operon gene map for each strain. To do so, we retrieved the raw DNA sequences for each sample from contigs built during our MAG assembly on KBase and extracted information such as sequence length, orientation, and estimated gene ontology. To investigate the phylogenetic relationship of each operon in reference to that of the reference strain *C. braakii* MiYa, we concatenated the DNA sequence of each mapped gene into a supermatrix before running a multiple alignment and constructing a phylogenetic tree using a maximum-likelihood neighbor-joining (NJ) method. Analysis and tree visualizations were performed using the *R* packages *ggtree* (version 3.12.0) and *gggenes* (version 0.5.1.9002).

### 2.8 Nutritional Composition Analysis

To assess the nutritional composition of the cabbage and carrot ferments, the concentrations of six different fat-soluble vitamins were measured: Vitamin A, Vitamin D_2_, Vitamin D_3_, Vitamin E, Vitamin K_1_, and Vitamin K_2_. Briefly, frozen carrot and cabbage samples were thawed slowly in the refrigerator at 4 . Approximately 200 mg of the sample was placed in a 2 mL centrifuge tube, resuspended, and ground in 440 μL of 0.025 M KOH for 5 minutes. Samples were then mixed in 100 μL of 4 M HCl, followed by 700 μL of n-hexane, and subjected to ultrasound sonication at 4 °C for 30 minutes. After a 30-minute downtime at 4 °C, the samples were centrifuged at 12,000 rpm for 15 minutes at the same temperature. The upper n-hexane layer, containing the vitamin fraction, was transferred to a new centrifuge tube and concentrated using roto-vap. Samples were redissolved in 200 μL of acetonitrile and centrifuged at 12,000 rpm for 15 minutes. The supernatant was then transferred to a new vial for LC-MS electrospray ionization analysis using an Acquity UPLC instrument (Waters, Millford, MA). MultiQuant software was used to integrate the peak counts, which were calculated according to the constructed standard curve. Final sample measurements were reported in nanograms of vitamin per gram of raw sample (ng/g).

## 3. Results & Discussion

### 3.1 Observations about Fermentation

As a starting point, we were interested in understanding how seasonal frost affects the culinary qualities of ferments. The idea is that seasonal frost should not negatively impact the usability for fermentation or the taste and texture of fermented cabbage and carrot. Such an impact would reduce the ferments’ commercial value and, thus, the attractiveness of frost exposure as a season-extension strategy. Confirming previous observations that had initially encouraged our investigation, we found that all crops fermented well, yielding products suitable for consumption. Crucially, we found that ferments from frost-exposed vegetables, or frost-conditioned ferments, outperformed traditional ferments. Frost-conditioned ferments were ready in 23 days, compared to 35 days for conventional ferments, or about 35% shorter completion time. Moreover, a panel of professional chefs at Blue Hill conducted a blind test of the different ferments. It concluded that frost-conditioned ferments presented a “brighter,” “less-muddied” acidity, and a “cleaner” flavor profile. In short, the frost-conditioned ferments were preferred to the traditional ferments made from crops harvested in the fall.

### 3.2 Characterizing the effect of frost on the microbial population of cabbage ferments

First, using *16S rRNA* amplicon sequencing, we characterized the different probiotic bacterial content of the eight cabbage ferments. We found a total of thirty-three genera, including *Latilactobacillus* sp., which accounted for 94.8 ± 42.3% of the total bacterial populations. The latter is a bacterium commonly associated with the fermentation of vegetables. Within the genus, *L. sakei* is one of best-known species and is particularly important in the production of sauerkraut, contributing to flavor development through the production of compounds such as ethanol, carbon dioxide, and volatile compounds ^51^.

We then compared the bacterial communities between frost-conditioned ferments and traditional ferments. We found that frost did not affect overall bacterial diversity in terms of the number of predicted bacterial types (*F*_(1,6)_ = 0.48; *p* = 0.52; **Figure 2A**) or relative abundance (*F*_(1,6)_ = 0.60; *p* = 0.47; **Figure 2B**). In other words, frost did not change the number of genera and their relative distribution in the ferments at the time of consumption. Yet, we found that frost had a weak but marginally significant effect on bacterial composition (permanova: *F*_(1,6)_ = 3.199; *p* = 0.04; **Figure 2C**). Interestingly, the only change of note was the presence of *Leuconostoc* sp. in three of the ferments initiated from frost-exposed plants, reaching a relative abundance of about 10% in two of the ferments. This finding is perhaps not surprising, given that *Leuconostoc* sp. is another common bacterium in fermented cabbage products, such as sauerkraut and, more specifically, kimchi ^52^. It is well-known for being psychrotolerant, or cold-tolerant, and able to grow at temperatures ranging from 2°C to 15°C without affecting fermentation ^53^. *Leuconostoc* sp. is often a dominant taxon found in kimchi fermentation, a practice prevalent in regions of Korea where cabbages and other ferments are exposed to colder temperatures ^54,55^.

**Figure 2.**
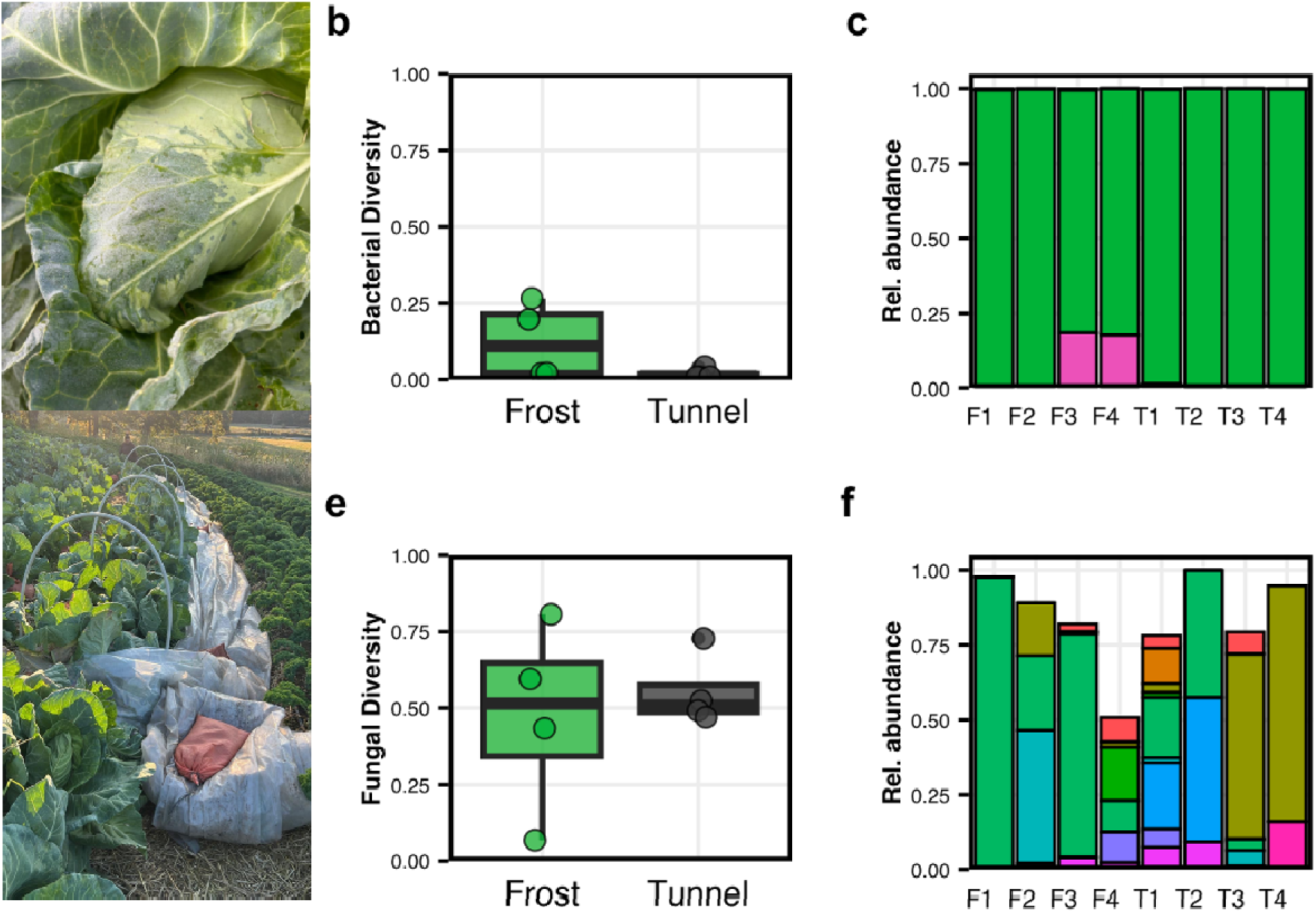
Effect of Frost on Microbial Communities Growing in Cabbage Ferments. In bacteria, we observed no difference in **a**) the number of predicted genera, and **b**) diversity measured as Pielou’s Evenness. However, we found a slight difference in **c**) overall population structures between the vegetables exposed to frost (green) and protected by tunnels (black). In fungi, we observed no difference in **a**) the number of predicted genera, and **b**) diversity measured as Pielou’s Evenness. However, we found a small difference in **c**) overall population structures between the vegetables exposed to frost (orange) and protected by tunnels (black). Each circle represents an estimate from a single ferment initiated from a specific plant. Each circle represents an estimate from one ferment initiated from a single plant. In bacteria, **c)** genera are identified as *Buttiauxella* (red), *Citrobacter* (blue), *Klebsiella* (green), *Latilactobacillus* (cyan), *Leuconostoc* (purple), *Pectobacterium* (gray), *Pseudocitrobacter* (brown), *Raoultella* (yellow), and *Yersinia* (orange). In fungi **f)** genera are identified as *Aspergillus* (orange), *Bullera* (brown), *Cladosporium* (khaki), *Cystofilobasidium* (gray), *Debaryomces* (dark green), *Penicillium* (teal), *Pseudogymnoascus* (blue), *Rhodotorula* (violet), *Saccharomyces* (purple), and *Trametes* (pink).

When considering fungal probiotics using *ITS2* amplicon sequencing, we found that communities varied widely between ferments, with *Dabaryomyces* sp. being the most common genus 64.8 ± 7.6%). Overall, we found that frost did not affect the number of different fungal genera in the ferments (*F*_(1,6)_ = 0.04; *p* = 0.85; **Figure 2D**). Interestingly, we found that frost had a weak but marginally significant effect on the distribution of the fungal species, tunnel ferments showing a higher level of diversity than ferments from outside cabbages (*F*_(1,6)_ = 4.89; *p* = 0.069; **Figure 2E**). While such changes did not contribute to an overall shift in composition between the two treatments (PERMANOVA: *F*(1,6) = 0.89; *p* = 0.64; **Figure 2F**), we found that single fungal ASVs exhibited significant changes between the treatments. More specifically, we found that *Lycoperdon* sp., *Saccharomyces* sp., *Humicola* sp., and *Fusarium* sp. were in higher abundance in frost-conditioned ferments, while *Penicillium* sp., *Chaetomium* sp., *Aspergillus* sp., *Rhodotorula* sp., and *Pseudogymnoascus* sp. were in higher abundance in tunnel ferments. While the role of fungi in lacto-fermentation is rarely considered, changes in fungal species could also affect the flavor profile of ferments.

### 3.3 Characterizing the effect of frost on the microbial population of carrot ferments

In fermented carrots, on the other hand, bacterial communities varied widely in composition, with *Citrobacter* sp. (39.8 ± 19.3%) being the most common genus, followed by *Raoultella* sp. (28.2 ± 16.0%) and *Latilactobacillus* sp. (11.5 ± 25.8%). While the role of *Latilactobacillus* sp. bacteria in food fermentation is extensively documented as discussed above, *Citrobacter* sp. and *Raoultella* sp. are usually considered opportunistic fermenters; present in ferments due to their ubiquity in soil environments ^56^. The potential for both bacteria in fermentation processes is being further considered as we expand our understanding of a broader range of fermented foods ^57–60^. For example, we observed, in a previous study, the presence of *Citrobacter* sp. in fermented evergreens associated with briny flavors, most likely explained by the production of volatile compounds ^36^. We also found *Yersinia* sp. in three of the carrot ferments; another bacterium not generally associated with healthy fermentation, and that will require further investigation.

Similarly to fermented cabbages, we found no significant differences in the number of general between frost-conditioned and tunnel-protected carrot ferments (*F*_(1,4)_ = 1.64; *p* = 0.25; **Figure 3A**) or in the diversity of bacterial type accounting for relative abundance of each type (*F*_(1,4)_ = 0.31; *p* = 0.61; **Figure 3B**). Here, however, we observed more significant changes in overall population structure between the two treatments (PERMANOVA: *F*_(1,6)_ = 4.71; *p* = 0.02; **Figure 3C**). More specifically, we found an increase in *Raoultella* sp. in frost-conditioned ferments. In contrast, we saw an increase in *Pseudomonas* sp. and *Pantoea* sp. in the ferments made from carrots protected from frost. Interestingly, there is no evidence that *Raoultella* sp. is cold-tolerant, especially when compared to *Pseudomonas* sp., which is documented to be cold-tolerant. This result is surprising to us, given the capacity of many *Pseudomonas* sp and *Pantoea* sp. species to infect plant tissue ^61–63^ we would have expected that frost-induced plant tissue damage would have offered further protection to the two bacterial genera ^64,65^. Given the rate of growth shown by common LAB in the frost-conditioned ferments, we suspect that it is the vigorous growth that reduces the abundance of the two opportunistic pathogens.

**Figure 3.**
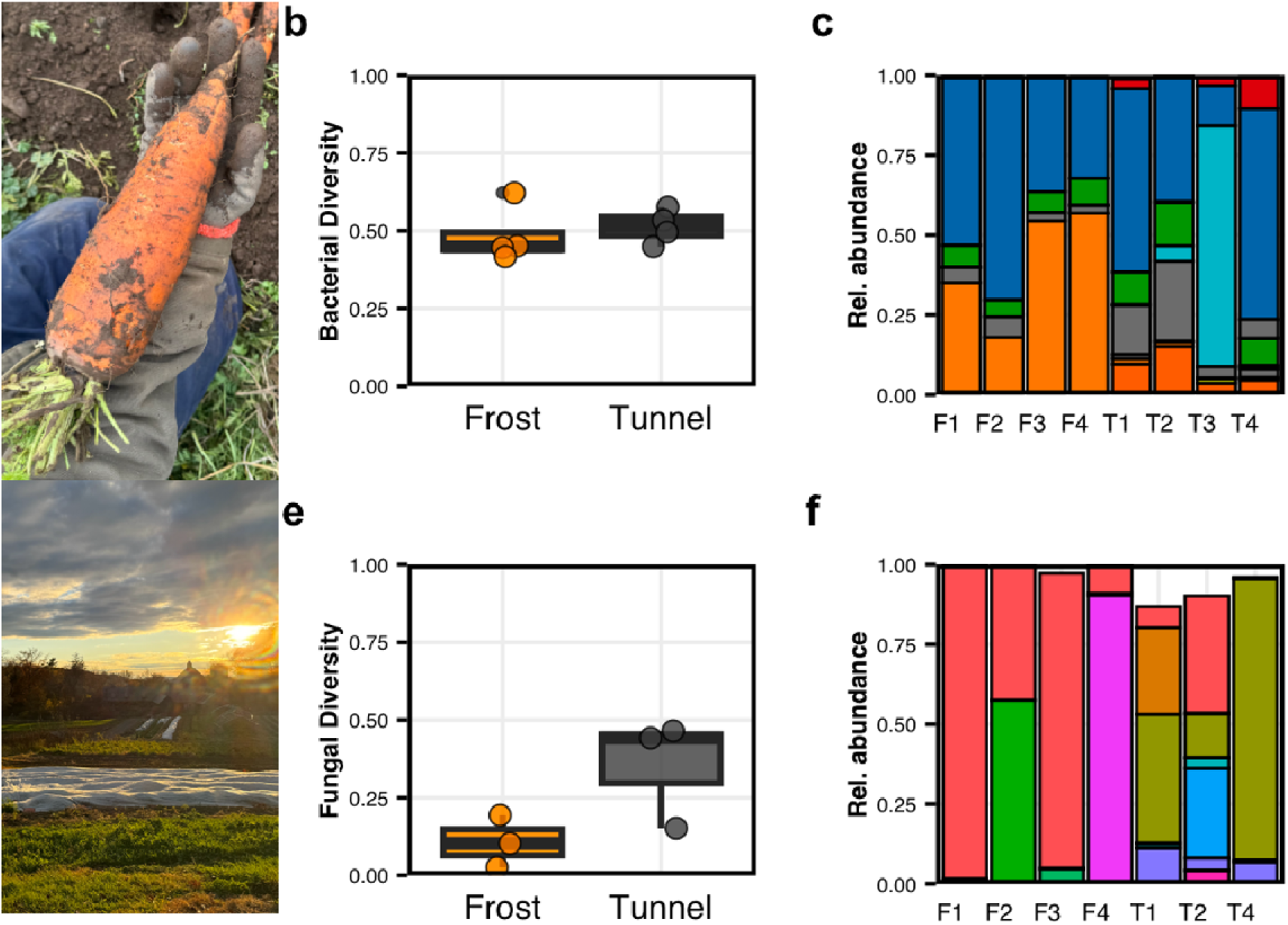
Effect of frost on microbial communities found in carrot ferments. In bacteria, we observed no difference in **a**) the number of predicted genera, and **b**) diversity measured as Pielou’s Evenness. However, we found a small difference in **c**) overall population structures between the vegetables exposed to frost (orange) and protected by tunnels (black). In fungi, we observed no difference in **a**) the number of predicted genera, and **b**) diversity measured as Pielou’s Evenness. However, we found a small difference in **c**) overall population structures between the vegetables exposed to frost (orange) and protected by tunnels (black). Each circle represents an estimate from one ferment initiated from a single plant. In **c)** genera are identified as *Buttiauxella* (red), *Citrobacter* (blue), *Klebsiella* (green), *Latilactobacillus* (cyan), *Leuconostoc* (purple), *Pectobacterium* (gray), *Pseudocitrobacter* (brown), *Raoultella* (yellow), and *Yersinia* (orange). In **f)** genera are identified as *Buttiauxella* (red), *Citrobacter* (blue), *Klebsiella* (green), *Latilactobacillus* (cyan), *Leuconostoc* (purple), *Pectobacterium* (gray), *Pseudocitrobacter* (brown), *Raoultella* (yellow), and *Yersinia* (orange).

Again, using *ITS* amplicon sequencing, we characterized the fungal populations found in fermented carrots. Once again, we found that Debaryomyces sp was the most common genus (39.2 ± 9.4%), followed by Fusarium sp. (16.4 ± 13.5%) and *Saccharomyces* sp. (13.2 ± 24.6%). Similarly to fermented cabbages, we found no significant differences in the number of genera between frost-conditioned and tunnel-protected ferments (*F*_(1,5)_ = 0.0004; *p* = 0.98; **Figure 3D**). Interestingly, as was the case in cabbage, we found that frost impacted the relative abundance of the fungal type, showing higher overall diversity in tunnel ferments (*F*_(1,5)_ = 7.62; *p* = 0.04; **Figure 3E**). While these changes in relative abundances, unlike those observed in bacterial populations, did not translate into significant changes in overall fungal composition (PERMANOVA: *F*_(1,5)_ = 2.30; *p* = 0.08; **Figure 3F**), we observed a few individual genera changing in abundance. The most important change is the increase in abundance of *Debaryomyces* sp. in the first-conditioned ferments. Indeed, particular species within the genus, such as *D*. *hansenii*, have been characterized as extremophilic, tolerating high salt and sugar concentrations as well as cold temperature ^66^. The genus is also often associated with the fermentation of meat products, such as sausages, where it is characterized as metabolically versatile, contributing significantly to flavor ^67^.

### 3.4 The effect of frost-conditioning on predicted metabolite profile

Using whole-genome sequencing to describe the overall microbial communities, we then investigate potential changes in metabolic profiles at the genomic level. In short, we identified all genes sequenced as part of the ferment communities and, based on their annotation, found a total of 393 out of 576 functional subsystems that contained annotated genes, between both groups. Out of the detected functional subsystems, cabbages had 244 detected subsystems with a total of 7,487 annotated genes; carrots had 386 subsystems with a total of 35,691 annotated genes (**Table S7**).

First, we searched for significant changes in any metabolic pathways that may exist between frost-conditioned ferments and tunnel-protected ferments. In cabbages, we identified seven metabolic pathways that exhibited significant changes (**Table S8**). In every case, but one, we found a metabolic increase in frost-conditioned ferments. The only exception was a reduction in acid resistance mechanisms, a pathway that is often associated with halotolerant bacteria or even foodborne pathogens. Examining metabolic pathways that increased in abundance, we identified four gene functions related to carbohydrate processing, such as D-ribose and fructose utilization, which were both more than three times more abundant in frost-conditioned ferments. We also found an increase in genes associated with threonine biosynthesis. This amino acid can enhance the flavor of food by adding sweetness and a richer, more layered taste, as well as contributing to the aroma and mouthfeel of food ^68^.

In carrot ferments, we identified a total of 26 metabolic pathways that exhibited significant changes in abundance between treatments (**Table S9**). From these, nineteen were in higher abundance in frost-conditioned ferments, including three gene functions related to carbohydrate utilization among the top three most considerable changes (i.e., rhamnose, fucose, as well as D-glucitol and D-sorbitol). Additionally, we identified genes related to the production of metabolites that could impact the flavor and nutritional properties of the ferments. For example, we observed a significant increase in the abundance of genes associated with coenzyme B_12_ biosynthesis, an essential vitamin primarily found in animal products and fermented foods ^69^. More specifically, the genes for the anaerobic biosynthesis pathway of cobalamin, or B_12_, were found in higher abundance in frost-conditioned ferments (**Figure 4A**), which confirms that fermented carrots can offer an option for increasing B_12_ availability.

**Figure 4.**
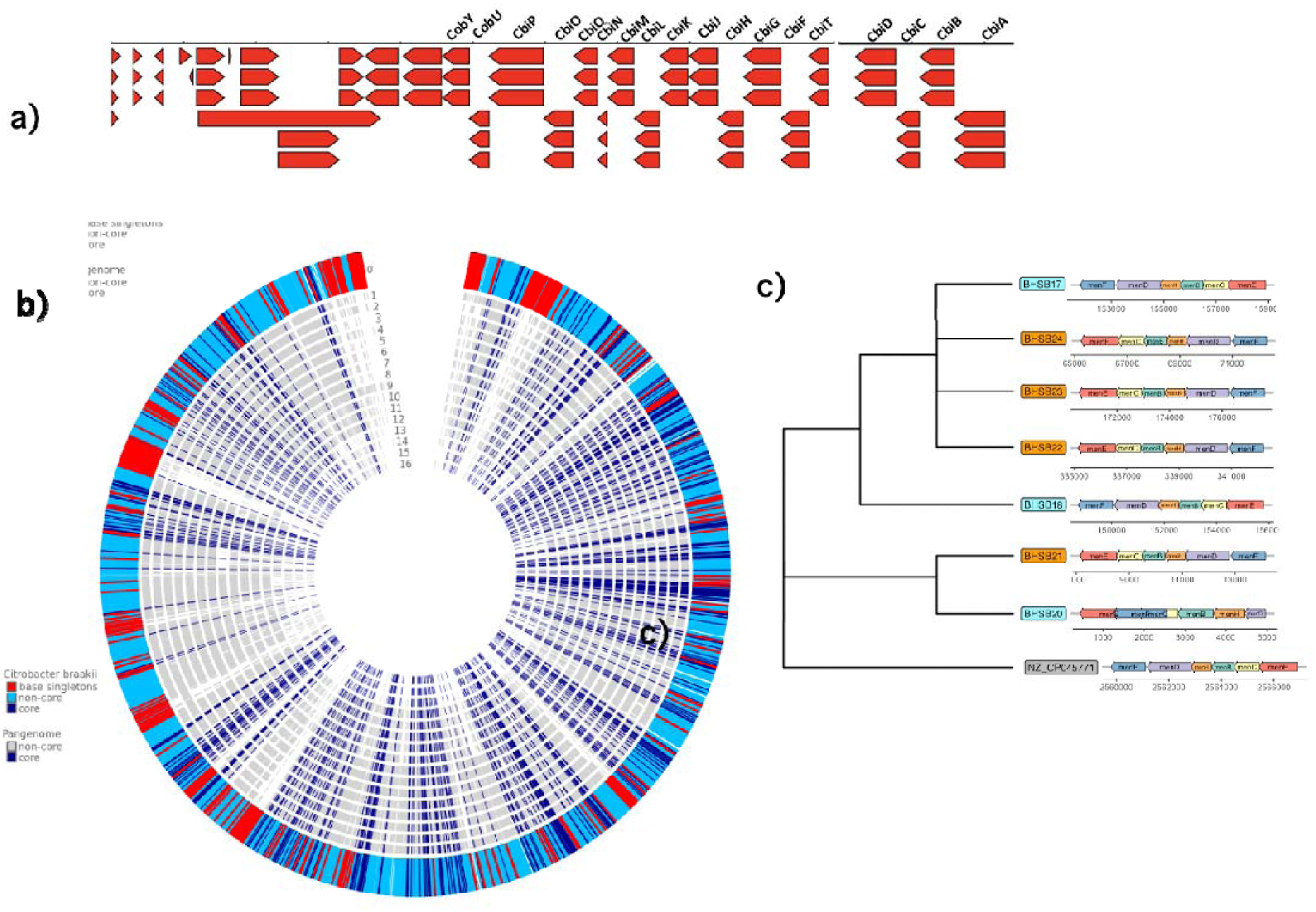
Distribution of annotated genes in cabbage and carrot ferments. **a**) Gene map and predicted mRNA and protein from a B_12_ vitamin biosynthesis pathway found in sample BHSB24. **b**) Total gene distribution in *Citrobacter braakii* genomes assembled from metagenomes from each sample. Alignment includes reference strain *C. braakii* MiYA. **c**) NJ-tree from multiple alignment of methionine operons found in C. braakii MAGs. Two genes are omitted from the gene map for visualization purposes.

We also found genes associated with the menaquinone pathway, responsible for the synthesis of vitamin K, in higher abundance in the frost-conditioned cabbage ferments. More specifically, we identified a cluster of genes, *men*F/*D*/H/B/C/E, associated with the intermediary reactions of chorismate to 1,4-dihydroxy-2-naphthoate and essential for producing menaquinol, in at least one of the genomes assembled from metagenomics data in every sample (**Table S10**). The genomes most closely matched to *Citrobacter braakii* MiYa, a strain known for producing menaquinone ^70^, and, crucially, showed a significant amount of variability in gene composition (**Figure 4B**). All strains exhibited some level of divergence from the reference genome, indicating that the *Citrobacter* species present in the ferments are likely unique to the environment studied.

When comparing the *men* operons across genomes, we found that each operon displayed a distinct gene arrangement, with at least one or two gene positions differing between samples. This further supports the placement of *C. braakii* MiYA as an outgroup (**Figure 4C**). Notably, the gene *memF* appeared consistently at the end of the operon in genomes from conventional ferments. In contrast, in both the reference genome and the frost-conditioned MAGs, *memF* was positioned among the first two loci. Interestingly, when examining the phylogenetic tree for each gene individually, we found that horizontal gene transfer could explain such modifications (**Figure S4**). Since genes located closer to the origin of replication can experience increased copy number during active cell division, this shift in positioning could enhance the expression of *memF* and, by extension, menaquinone (vitamin K) production. Given vitamin K’s role in bacterial anaerobic respiration, such a genomic arrangement may provide a selective advantage in the altered redox environment of frost-damaged vegetables.

Finally, although our results suggest that frost exposure may enhance metabolic potential in vegetable ferments, we considered the possibility that the observed increase in complete metabolic pathways could be an artifact of reduced contig fragmentation in the frost-conditioned samples. To account for this potential bias, we performed a BLAST-based validation to assess whether the relevant genes were located on full-length contigs. Using the reference genome *Citrobacter braakii* MiYA, we found that the gene sequences of interest consistently aligned with this species. In both treatments, we confirmed that the majority of genes involved in cobalamin and menaquinone biosynthesis were located on contiguous sequences with high similarity to the reference genome, supporting the robustness of our pathway-level comparisons.

### 3.5 Characterizing the effect of frost on ferment’s nutritional qualities

We then investigated the possible effect of frost on the nutritional content of our ferments. Here, we focused our attention on fat-soluble vitamins, namely vitamins A, E, K_1_, and K_2_, which are expected to be present in both cabbages and carrots. More specifically, carrots are recognized as an excellent source of beta-carotene and vitamin A, as well as vitamin E ^71^. On the other hand, cabbage is known to be an important source of Vitamin K_1_ and K_2_ and contains some vitamin E ^72^. Finally, fat-soluble vitamins are also known to be more stable during food processing, such as frying and fermentation ^73–75^, enabling us to test whether frost would impact the presence of essential nutrients expected to be found in ferments.

To validate our methodology, we first compared nutritional qualities between fermented cabbage and fermented carrots (**Figure S2**). As expected, our results show that Vitamin A (**Figure S2A**) and Vitamin E are enriched in carrots compared to cabbages (Figure S2B). We also found, as expected, that Vitamin K_1_ was higher in cabbage compared to carrots (**Figure S2C**), but there was no difference in Vitamin K_2_ (**Figure S2D**). The abundance of Vitamin K_1_ in cabbage is explained by the role phylloquinone plays in photosynthesis in leafy greens. Lastly, given that neither cabbage nor carrot contains menaquinone (vitamin K_2_), the presence of this nutrient in all ferments confirms the vital role of fermentation in enhancing nutritional values and supporting a healthy diet.

Overall, we found no negative impact of frost on nutritional value. In itself, this is a significant observation that demonstrates that frost had no detrimental effects on the availability of these vitamins in two important crops commonly used for fermentation. Looking into cabbages, to frost and the fermented cabbage protected from the frost: Vitamin A (*F*_(1,6)_ = 3.11; *p* = 0.13; **Figure 5A**), Vitamin E (*F*_(1,6)_ = 1.25; *p* = 0.30; **Figure 5B**), Vitamin K_1_ (*F*_(1,6)_ = 0.03; *p* = 0.87; **Figure 5C**), and Vitamin K_2_ (*F*_(1,6)_ = 1.47; *p* = 0.27; **Figure 5D**). Similarly, we saw no difference in the availability of Vitamin A (*F*_(1,6)_ = 1.62; *p* = 0.48; **Figure 5E**) and Vitamin K_1_ (*F*_(1,6)_ = 0.03; *p* = 0.87; **Figure 5F**) in frost-conditioned carrot ferments and traditional ferments. Crucially, however, we observed increases in Vitamin E (*F*_(1,6)_ = 23.35; *p* = 0.003; **Figure 5G**) and Vitamin K_2_ (*F*_(1,6)_ = 11.66; *p* = 0.01; **Figure 5H**).

**Figure 5.**
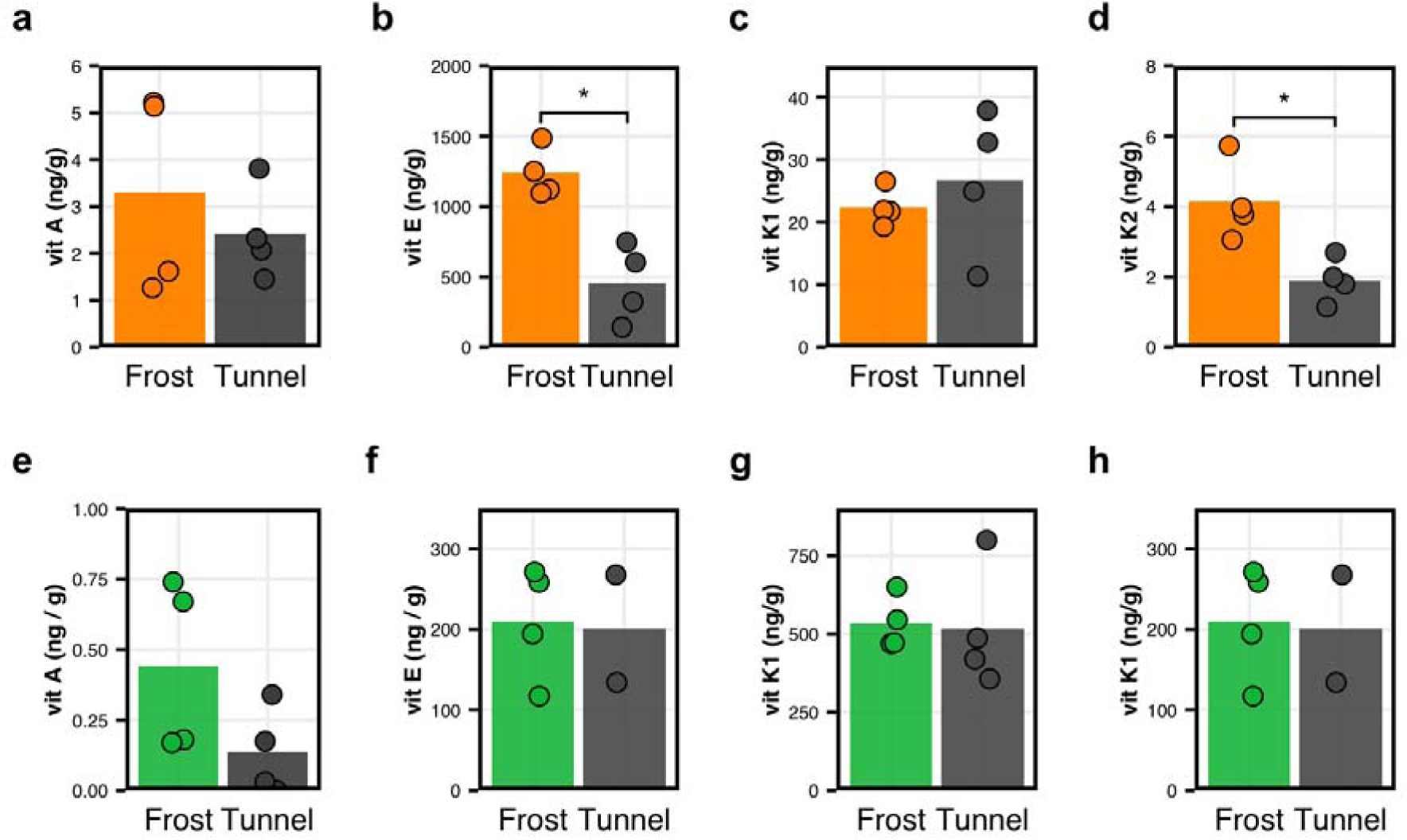
Effect of frost on nutrient bioavailability in fermented cabbage. For each gram of fermented food product, the concentration **a)** Vitamin A; **b)** Vitamin D_2_; **c)** Vitamin D_2_, **d)** Vitamin E, **e)** Vitamin K_1_, and **f)** Vitamin K_2_ was calculated in nanograms. Box plots show the median as well as the interquartile range. Individual points represent values for each sample. Significant differences, estimated using the Kruskal-Wallis test, are shown as “*” for *p* < 0.05.

The increase in vitamin content in carrots could be explained by the direct effect of frost on the crop before fermentation, the impact of frost on the microbial communities present on the crop at the time of exposure, which subsequently shape fermentation, or a combination of the two. Indeed, frost can affect the structure of cell walls, potentially allowing for better release of these vitamins during fermentation. Frost is also known to stress plants and alter their metabolic pathways, which may influence the content of specific nutrients. In some cases, this could lead to increased production of protective compounds, including antioxidants and vitamins, as a plant’s response to stress. Finally, carrots and cabbage, when exposed to frost, may experience changes in their lipid metabolism or the biosynthesis of fat-soluble vitamins, such as vitamin K and vitamin A precursors. In the context of our study, however, only the first scenario is likely to explain, at least partly, our results, given the short exposure to frost.

Finally, fermentation is also known to increase the bioavailability of certain phytochemicals ^34^. For example, as discussed above, vitamin K_2_ is not initially present in raw plant materials; instead, it is produced by bacteria during fermentation. This suggests that, at least in the context of vitamin K_2_, the effect of frost on menaquinone was modulated during the fermentation process. Vitamin K_2_ is generally biosynthesized during the later stage of fermentation when *Lactobacillus* sp. ferments available sugars ^76^. We believe that frost either selected for strains of *Lactobacillus* sp. that produced elevated levels of K_2_ or, perhaps, accelerated fermentation by increasing the availability of sugars following exposure to frost. Given that we observed fermentation to be faster in frost-conditioned ferments, we can speculate that the latter had more time to accumulate vitamin K_2_, thereby adding to the nutritional benefits of frost exposure.

## 4. Conclusion

Farming in cold climates, such as New York and the American Northeast more generally, presents a unique set of challenges due to the shorter growing season ^21,22^. Further exacerbated by climate change, rapid shifts in extreme weather conditions have increased in frequency over the past decade ^23^. The use of hardier crops such as kale, Brussels sprouts, cabbage, and carrots, which show tolerance to colder weather, provides farmers with the opportunity to extend the growing season in the field and at the market. Yet, such a season extension strategy increases the risk of crops being exposed to a frost event. While frost is known to affect the flavor profile of certain vegetables, such as carrots, the impact of frost on the use of these vegetables in fermentation remains to be tested.

Here, we demonstrate that a single seasonal frost event not only does not negatively impact the quality of fermented cabbages and carrots, but can increase the nutritional value of these ferments. More specifically, using microbiological and biochemical analyses to compare ferments made from vegetables exposed to frost with those grown under similar conditions but protected from frost, we demonstrate that a single environmental frost event did not significantly alter the microbial communities that developed during fermentation, nor did it reduce the bioavailability of key fat-soluble vitamins. On the contrary, we found that frost exposure increased the bioavailability of vitamin A and vitamin E in fermented carrots, as well as vitamin K□ in fermented cabbage. Thus, our findings suggest that frost exposure can enhance not only the sensory appeal but also the health-promoting potential of fermented vegetables.

Our study highlights an opportunity for farmers and food producers in the Northeastern US to differentiate their cabbage and carrots from those grown in warmer climates, such as California. By leveraging the region’s distinctive climatic conditions, particularly exposure to seasonal frost, Northeastern farmers can produce vegetables perceived as more flavorful and potentially more nutrient-dense. With consumers increasingly seeking high-quality, flavorful, and nutrient-rich foods, the elevated levels of vitamins A, E, and K in frost-exposed, fermented carrots and cabbage may appeal to health-conscious buyers willing to pay a premium.

In addition, cultivating hardy crops in colder climates presents an opportunity to reduce the environmental costs of agriculture, particularly by lowering the need for irrigation and synthetic fertilizers. While large-scale cabbage production in arid regions, such as California’s Russian Valley, is currently more water-efficient per unit of yield than production in colder areas like New York’s Hudson Valley, the latter could become even more water-efficient on an annual basis if season extension practices are adopted.

Moreover, higher atmospheric temperatures are predicted to increase soil evaporation and the need for crop irrigation, particularly in arid climates ^11^. Ultimately, conventional efficiency metrics focus solely on crop weight; based on our findings, it is plausible that colder climates could yield greater nutritional returns per gallon of water used ^77^. This observation suggests a potential case for indexing water use in terms of nutritional or economic value. Although speculative, such considerations align with emerging global efforts to relocate crop production in ways that reduce agriculture’s overall environmental impact ^5^.

At the molecular level, our results suggest that changes in microbial populations resulting from frost exposure are, at least in part, responsible for the increase in nutritional value. As freezing and thawing alter the physicochemical landscape of plant tissues and create a hostile environment ^78^, one possible mechanism that could offer some protection for bacteria in such an environment is the enhanced production of vitamin K (menaquinone), a key cofactor in bacterial electron transport under anaerobic conditions ^79,80^. Interestingly, we found an increase in the copy number of the menaquinone biosynthesis pathway in bacterial genomes isolated from frost-conditioned samples. In addition, we observed an instance where a core gene in the menaquinone biosynthesis pathway, i.e., *memF*, was positioned closer to the origin of replication in a genome isolated from frost-conditioned samples, which is predicted to increase gene expression overall by ^81%^. In this context, strains with higher menaquinone production may be better suited to navigate the altered redox balance during the early stage of fermentation in post-frost samples. Similarly, we identified an increase in genes associated with propionate production, an organic acid, in microbial genomes isolated from frost-conditioned carrot ferments. Interestingly, propionate can impact the sharpness of the ferments ^82^ and their health benefits by acting as a prebiotic, promoting the growth of beneficial gut microbiota^83,84^.

It is essential to note, however, that our study does not provide a comprehensive picture of the process by which frost exposure affects nutrient bioavailability. For one, the variability in fermentation duration between the different ferments, which was necessary to attain the desired flavor profile, could have influenced the results. For example, longer fermentation could result in higher nutrient bioavailability. Yet, our findings show that the levels of vitamin K_2_, a chemical solely produced by the activity of fermenting bacteria ^70,85^, were similar in all ferments, suggesting that this is likely not the case. Additionally, as discussed above, our study does not enable us to fully understand whether the effect of frost exposure on ferments results from impacts on plant structure and physiology or impacts on the microbial populations inhabiting the plants. While our study suggests that both mechanisms are at play, future work will measure changes in a broader range of phytochemicals and will include raw vegetables immediately after frost exposure as well as after a set time for fermentation.

In conclusion, our study supports the idea that producing cabbage and carrots in the Northeast results in more flavorful and potentially more nutritious ferments. Extending the growing season into the fall and beyond the first seasonal frost can not only be part of a suite of practices that reduce the environmental costs of agriculture, but also contribute to creating a more desirable product. This latter point is crucial in developing incentives for farmers, as programs tied to short-term benefits tend to have higher adoption rates than those aimed solely at delivering ecological services ^86^. In short, by emphasizing these environmental advantages, Northeast farmers can build a distinct brand identity and enhance their economic returns. At the same time, offer a unique and desirable food product rooted in the region’s terroir and embedded in a more sustainable, resilient agricultural approach.

## Acknowledgements

This work was funded by the Stone Barns Center for Food and Agriculture (Pocantico Hills, NY, USA) and Bard College (Annandale-on-Hudson, NY, USA). This work was also supported in part through the NYU IT High Performance Computing resources, services, and staff expertise (New York, NY, USA). The authors would like to thank Daniel Bartush and Shannon Ryan for their assistance in the field and the lab, respectively, Susi Lu and Kamryn McKenna for their help with the vegetables illustrations in Figure 1, as well as Ani Alpert and Pia Sörensen for their comments and suggestions on the manuscript.

## Contributions

The study was designed by AL, JG, DB, SJ, and GGP. AL and JG collected samples. SJ and GGP processed samples and analyzed data. AL, JG, SJ, and GGP wrote the manuscript. SJ and GGP made final revisions on the manuscript.

## Data availability

Raw data (i.e., fastq files) of *16S rRNA* and *ITS* amplicon sequencing and whole-genome sequencing are available on the NCBI’s SRA database using accession number PRJNA565393. Other data and R scripts for data analysis are included as supplementary materials.

## Competing interests

The authors declare that they have no conflicts of interest. JG, farm manager at Stone Barns Center for Food and Agriculture, as well as AL and DB, both associated with the same center, are involved in the sale of farm products; however, this is not a primary objective of the organization. All authors, including corresponding author GGP, were free to write and conclude the paper independently, without influence from any of the organizations involved.

## APPENDIX

**Table S1. Weather data for growing season 2023, Harrison, NY.**

**Table S2. List of all ASVs identified in *16S rRNA* amplicon data.**

**Table S3. List of all ASVs identified in *ITS* amplicon data.**

**Table S4. List of all taxa identified in *16S rRNA* amplicon data.**

**Table S5. List of all taxa identified in *ITS* amplicon data.**

**Table S6. List of all samples and associated metadata.**

**Table S7. List of all predicted genes identified from whole-genome sequencing data.**

**Table S8. List of all metabolites that changed in frequency in cabbage ferments.**

**Table S9. List of all metabolites that changed in frequency in carrot ferments.**

**Table S10. Descriptive statistics associated with metagenomes-assembled genomes.**

---

**Figure S1.**
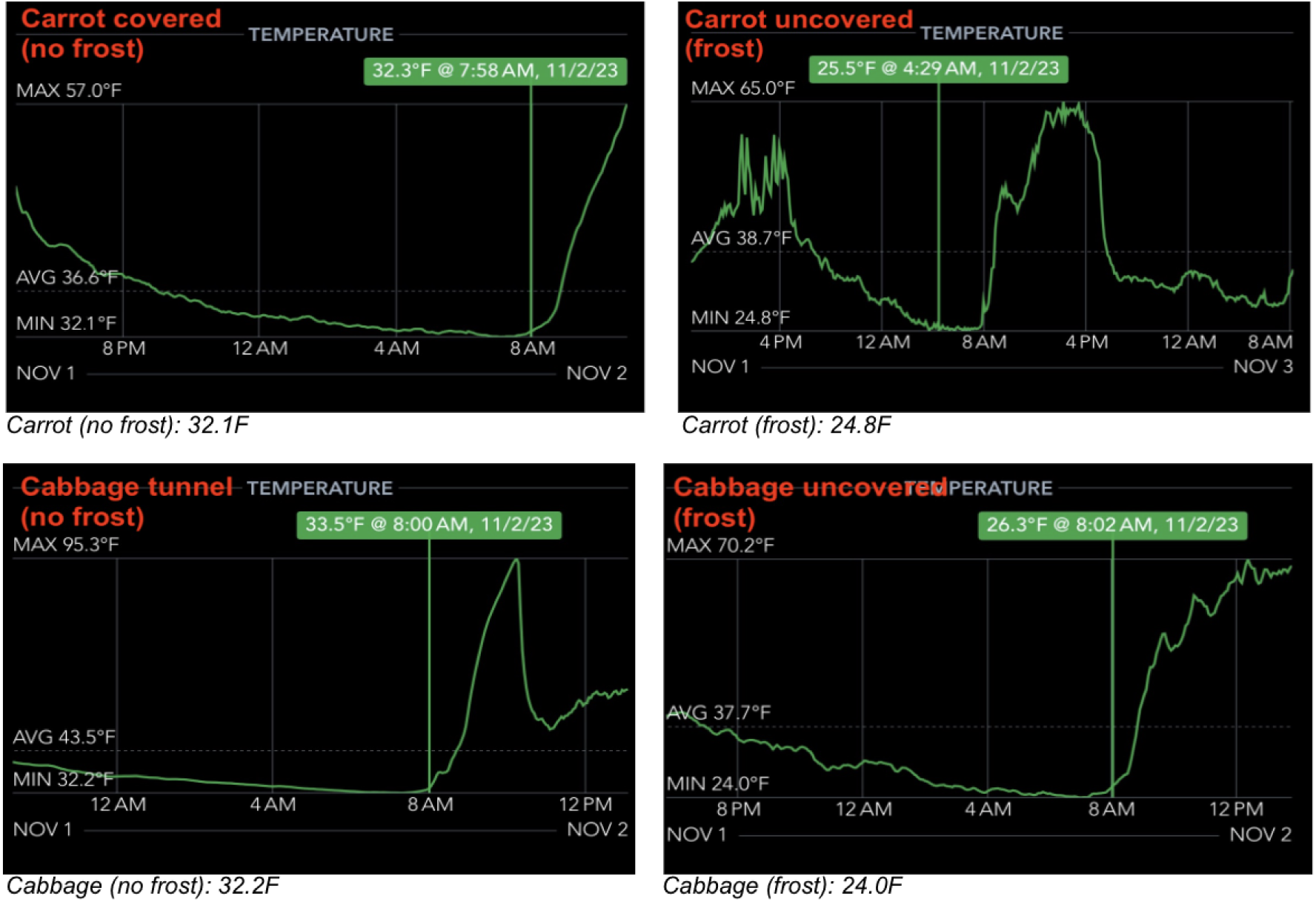
Recording of Air Temperature on November 1, 2023. Temperature (LJ) was measured using air sensor probes (SensorPush, Brooklyn, NY) at 10 cm above ground in **a)** covered carrots; **b)** uncovered carrots, **c)** covered cabbage; and **d)** uncovered cabbage.

**Figure S2.**
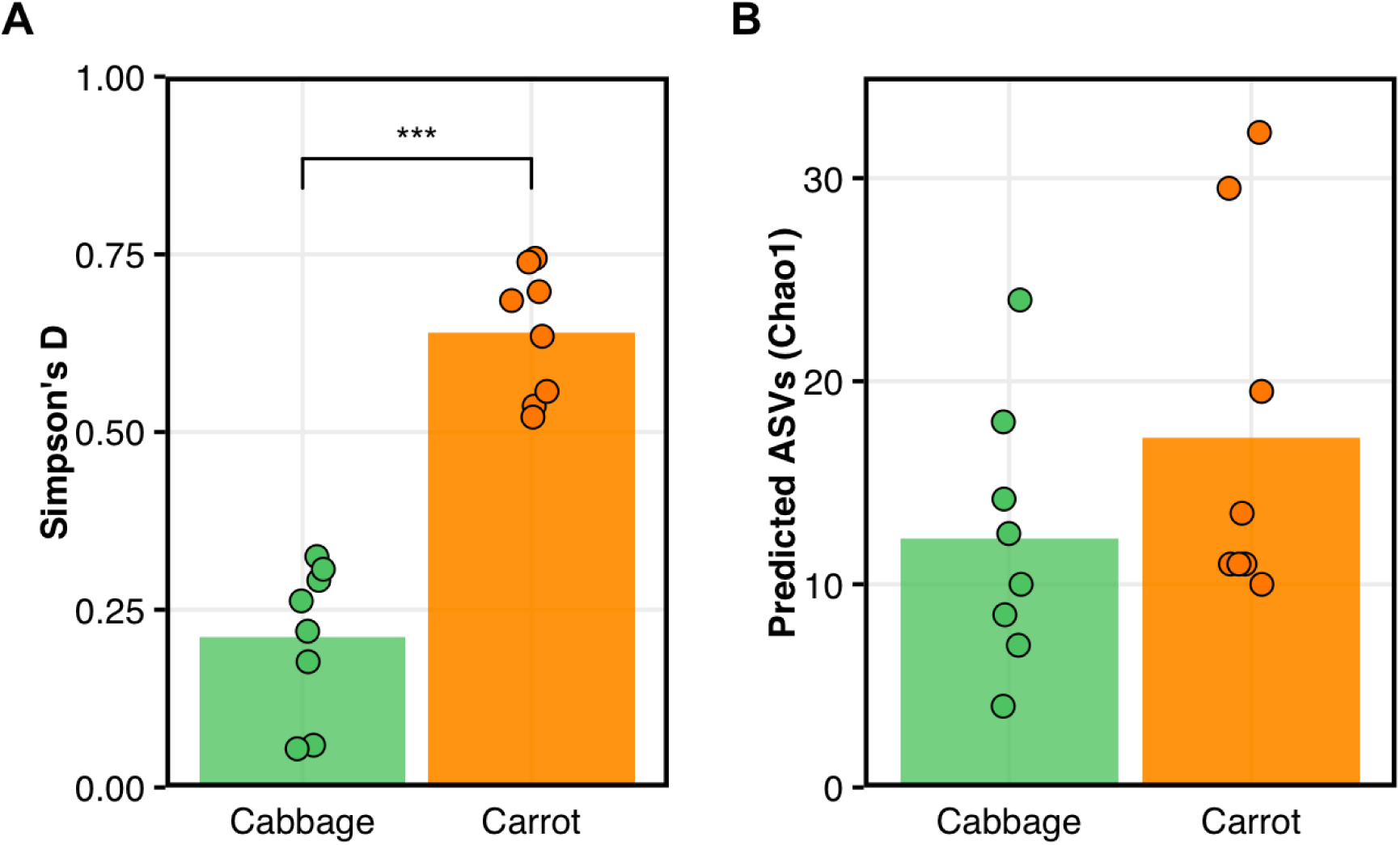
Bacterial Diversity in Cabbage and Carrot Ferments. **a**) Carrot ferments (vermillion) showed higher diversity measured as Simpson’s *D* than cabbage ferments (green) (STATS), while **b**) there was no difference in the number of predicted ASVs (STATS). Each circle represents an estimate from one ferment initiated from a single plant, and the top of the bars represents the mean for each group.

**Figure S3.**
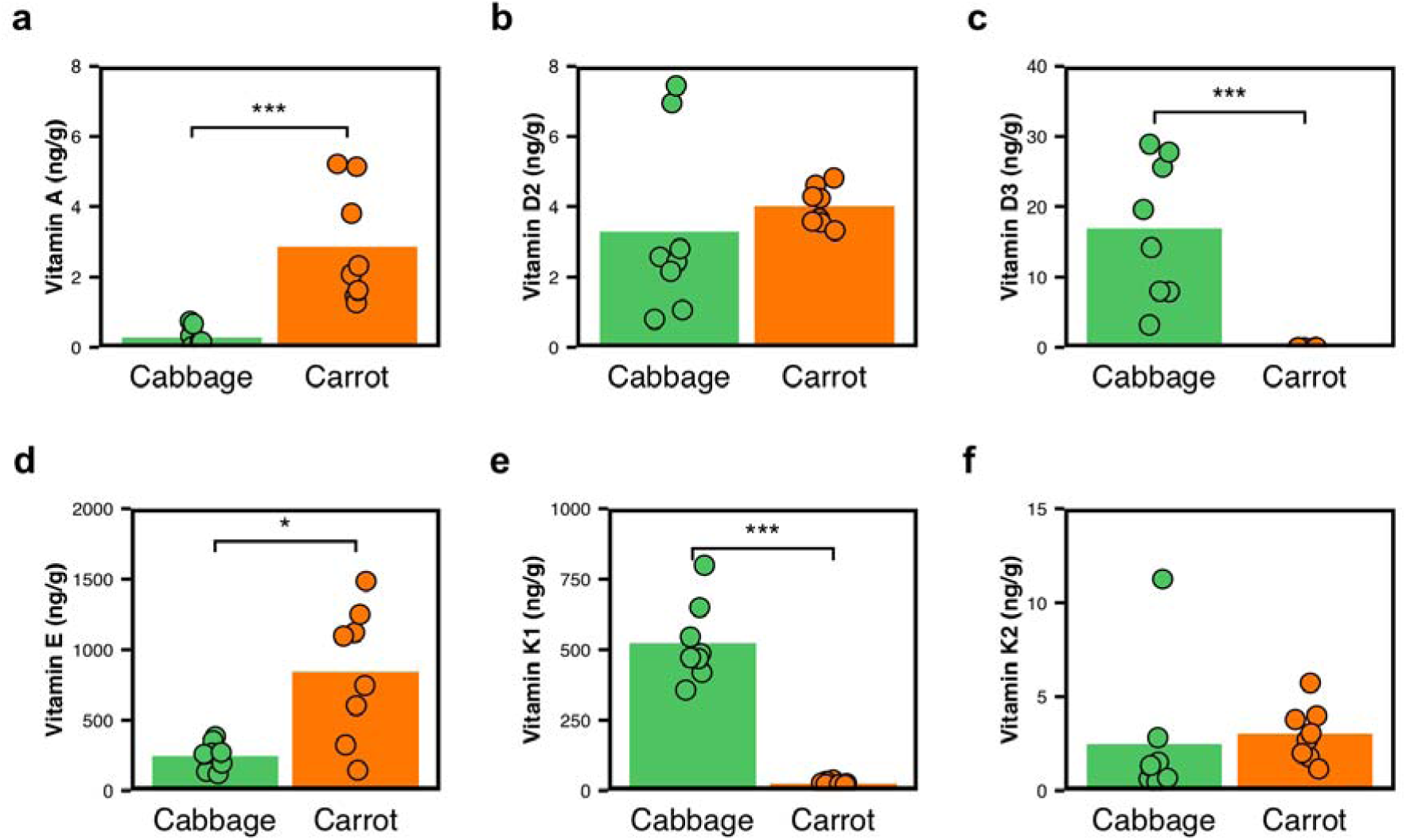
Nutritional Qualities of Fat-Soluble Vitamins in Fermented Cabbages and Carrots. For each gram of fermented food product, the concentration **a)** Vitamin A; **b)** Vitamin E; **c)** Vitamin K_1_; and **d)** Vitamin K_2_ was calculated in nanograms. Box plots show the median as well as the interquartile range. Individual points represent values for each sample. Significant differences, estimated using the Kruskal-Wallis test, are shown as “*” for *P* < 0.05.

**Figure S4.**
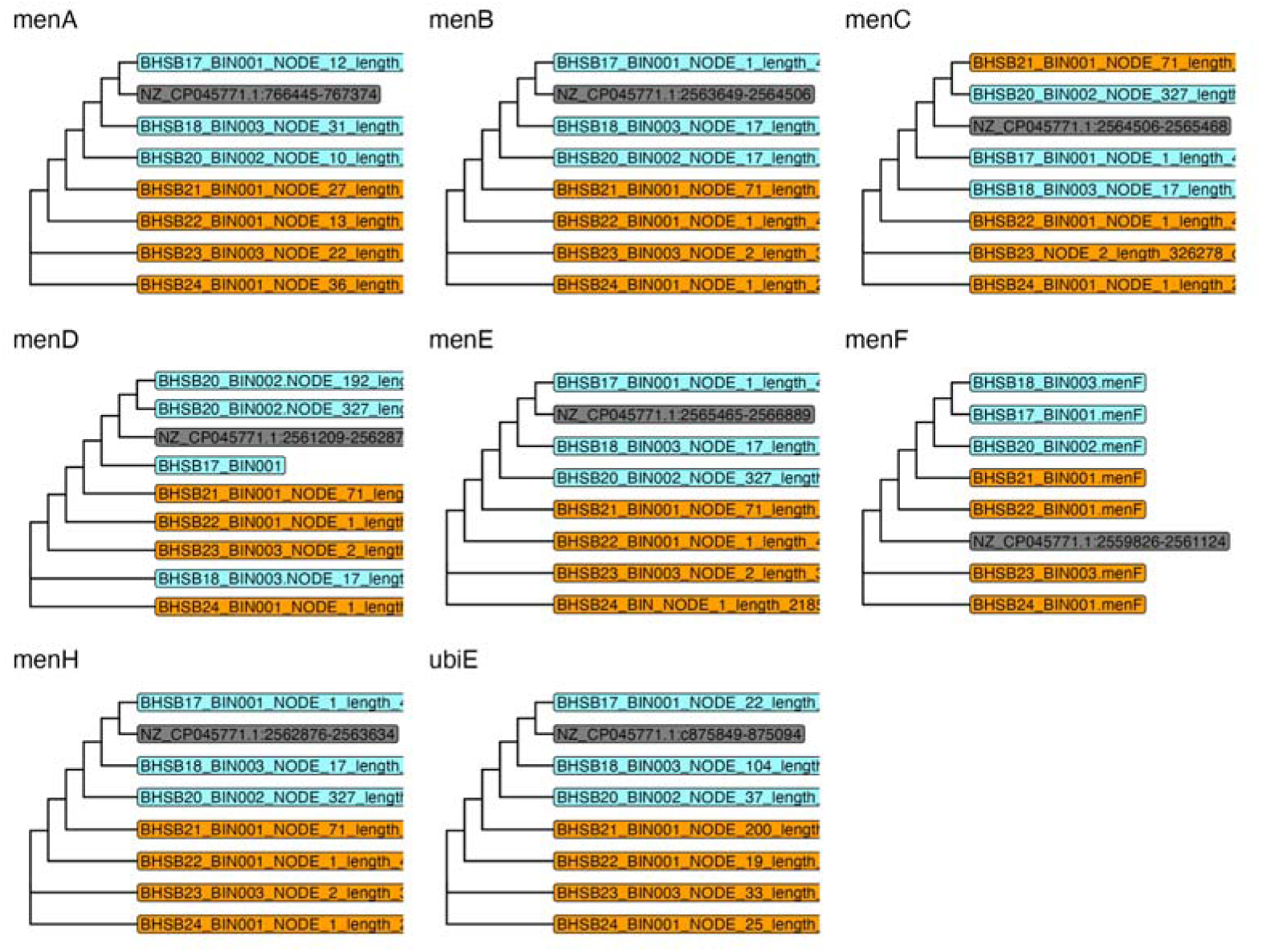
Phylogeny of Individual Genes Identified Within the *mem* Operon. NJ trees were constructed from multiple alignments run for each gene found in the men operon mapped on metagenome-assembled genomes found in each sample. *Citrobacter braakii* MiYa was used as a reference strain.

## Notes

https://www.ncbi.nlm.nih.gov/bioproject/?term=PRJNA1281670

