## Supplementary material for "Seasonal frost improves probiotic and nutrient availability in fermented vegetables": Table S1

| **Table 1. Weather data for growing season 2023, Harrison, NY.** | | | | | | | | | | | |
| --- | --- | --- | --- | --- | --- | --- | --- | --- | --- | --- | --- |
|  | **Precipitation (mm)** | | |  | **Max temp. (℃)** | | |  | **Min temp. (℃)** | | |
| **Month** | *2023* | *AVG* | *SD* |  | *2023* | *AVG* | *SD* |  | *2023* | *AVG* | *SD* |
| **July** | 0.34 | 3.2 | 1.8 |  | 75.93 | 82.9 | 3.0 |  | 58.27 | 66.2 | 3.0 |
| **August** | 4.78 | 3.3 | 2.6 |  | 78.45 | 81.7 | 2.2 |  | 78.45 | 67.6 | 4.2 |
| **September** | 5 | 3.8 | 3.1 |  | 74.33 | 75.2 | 1.8 |  | 60.1 | 58.8 | 2.1 |
| **October** | 3.96 | 3.3 | 2.5 |  | 65.61 | 64.5 | 2.9 |  | 50.19 | 47.3 | 3.5 |
| ***Stats.**** | *14.08* | *13.6* | *5.0* |  | *73.58* | *76.1* | *1.2* |  | *61.8* | *60.0* | *1.2* |
| **Descriptive statistics: total precip. for precipitation and average for temperature.* | | | | | | | | | | | |
